# The molecular basis of the anticancer effect of statins

**DOI:** 10.1101/2024.02.05.578869

**Authors:** Giovanni Buccioli, Carolina Testa, Emanuela Jacchetti, Pietro Pinoli, Stephana Carelli, Stefano Ceri, Manuela T. Raimondi

## Abstract

Statins, one of the most used class of cardiovascular drugs with the primary function of reducing blood cholesterol levels, exert their effect by inhibiting the enzyme HMG-CoA reductase, the key player in cholesterol biosynthesis. While the primary indication for statins is the prevention of cardiovascular diseases, there has been growing interest in their potential anticancer effects. However, the current evidence on these effects is largely based on epidemiological observations and preclinical research, not yet substantiated by knowledge of the mechanisms behind it. Here we show that statins have an anticancer effect as they exploit the principle of Synthetic Lethality, a concept in which the combination of two non-lethal genetic or molecular events results in cell death or impairment. When either of these events occurs alone, it is not lethal, but when they happen coupled, they create a lethal condition for the cell. In this work we report that statins emerged from a computational data analysis that we performed on approximately 37,000 synthetic lethality couples. We performed this analysis to select repurposable drugs that could target genes involved in Synthetic Lethality couples with metastatic genes. We validated our discovery *in vitro* by drug tests performed on cell lines derived from cancers of the breast, ovary, and cervix. Our data-driven drug repurposing strategy allowed us to understand the molecular basis of the anticancer effect of statins, a discovery which can be directly translated into practical clinical applications in oncology.

## 1 Introduction

Cancer is an astonishingly complex disease and continues to challenge medical research and therapeutic strategies, with 1,918,030 new cases and 609,360 cancer related deaths in 2022 [1]. In spite of the considerable progresses achieved in cancer research, traditional treatments are still accompanied by well-known side effects and various drawbacks. For instance, surgery and radiotherapy, beyond their positive effects, often inadvertently accelerate tumor growth and invasion due to the body’s inflammatory response and the variable cellular susceptibility to radiation [2, 3]. Meanwhile, cytostatic chemotherapy, while effective in reducing primary tumor volume [4], is frequently undermined by the emergence of resistance mutations in cancer cells [5]. This resistance often leads to the progression of tumors with the development of more aggressive phenotype [6].

Metastatic tumors, due to their diffuse localization and acquired resistance to cytostatic agents, are particularly challenging to treat. Despite the availability of over 200 approved anti-cancer drugs, none have proven effective in inhibiting or treating cancer metastasis [7], underscoring the urgent need for innovative therapeutic regimens. In this scenario, immunotherapy has emerged as a promising approach, leveraging the body’s immune system to target cancer cells [8]. However, its efficacy in treating solid tumors has been limited due to their inherent complexity [9]. Furthermore, recent studies have identified instances of secondary malignant tumors following therapy with CAR-T cells [10].

The newly emerging therapeutic approach based on Synthetic Lethality (SL) could offer an encouraging solution [11]. It hinges on the simultaneous suppression of two genes, leading to cellular lethality, while the inhibition of each gene in isolation remains a non-lethal event [12]. It thus selectively targets cancer cells with specific mutations, sparing healthy cells and reducing toxicity [11]. This approach could be particularly potent for metastatic tumors, which tend to harbor more genetic mutations [13], thereby potentially exerting anti-metastatic and migrastatic effects, when treating secondary tumors and preventing their formation. SL therapy, therefore, holds the promise of providing a more effective, personalized treatment with fewer side effects [14].

SL therapeutics, despite their promise, have encountered obstacles in clinical translation [15]. The challenges are twofold: not only is the laboratory-identification of robust SL gene pairs from the myriad possible combinations in a mammalian cell a complex task [15], but the subsequent formulation and testing of a drug that effectively targets these identified genes also requires significant research investment [16]. Drug repositioning, the practice of repurposing existing drugs for new therapeutic applications, emerges as a promising strategy in the challenging landscape of drug development [17]. It offers reduced development timelines and financial burdens compared to traditional drug development [17]. Drug repurposing can be experimental-driven, often arising serendipitously [18], or data-driven, a hypothesis-driven approach that uses big data to identify drugs against targets [19]. The latter transforms system biology data into predictions of druggable targets, ideally providing an FDA-approved compound with potential modulatory functions [19]. This approach requires accurate computational pipelines and algorithms for data integration and has been facilitated by the accumulation of high-throughput data and advances in computational and data sciences. In these circumstances, thanks to computer-aided approaches, it is possible to turn non-targeted (old) therapies into personalized treatments by selecting better responders’ patients [20]. Currently, there are approximately 2,500 drugs that have received approval from the FDA [21]. Regrettably, a mere three drugs have been repurposed for cancer treatment: the Bacillus Calmette–Guerin vaccine for superficial bladder cancer, thalidomide for multiple myeloma, and propranolol for infantile hemangioma [22]. Furthermore, in the current scientific landscape, there is a stark absence of research that utilizes SL for the repositioning of drugs with antimetastatic and migrastatic effects. This glaring omission not only underscores the urgency for innovative research but also highlights the potential for a paradigm shift in the treatment of metastatic diseases. The exploration of this uncharted territory could potentially herald a new era in cancer therapy, revolutionizing our approach and transforming patient outcomes. Our research, integrating both experimental and computational studies, is aimed at developing a comprehensive approach to investigate repurposable drugs for the treatment of metastatic solid tumors, leveraging the concept of SL. This is achieved through the creation of integrated databases and computational analysis of diverse datasets, with the goal of identifying the most promising candidates for experimental validation. Among the top candidates, we selected Simvastatin, a member of the statin family, due to its widespread prescription [23] and retrospective meta-analyses [24, 25] highlighting its antitumor effect. Furthermore, we explore the potential of combining repurposed drugs with anti-metastatic effects and chemotherapeutics with cytostatic effects, potentially leading to therapies with lower cytotoxicity but enhanced efficacy. For instance, Simvastatin exhibits strong synergy with Temozolomide. Our work represents a pioneering effort in integrating computational and experimental approaches to investigate repurposable drugs with migrastatic and anti-metastatic effects on metastatic solid tumors, leveraging the potential of SL, and ultimately demonstrating the antitumor therapeutic principle of statins.

## 2 Results

### 2.1 Data-driven identification of the best repurposable candidates

The comprehensive framework of our study, as illustrated in Figure 1, encapsulates the methodology employed to discover repurposable drugs that inhibit genes forming

**Fig. 1:**
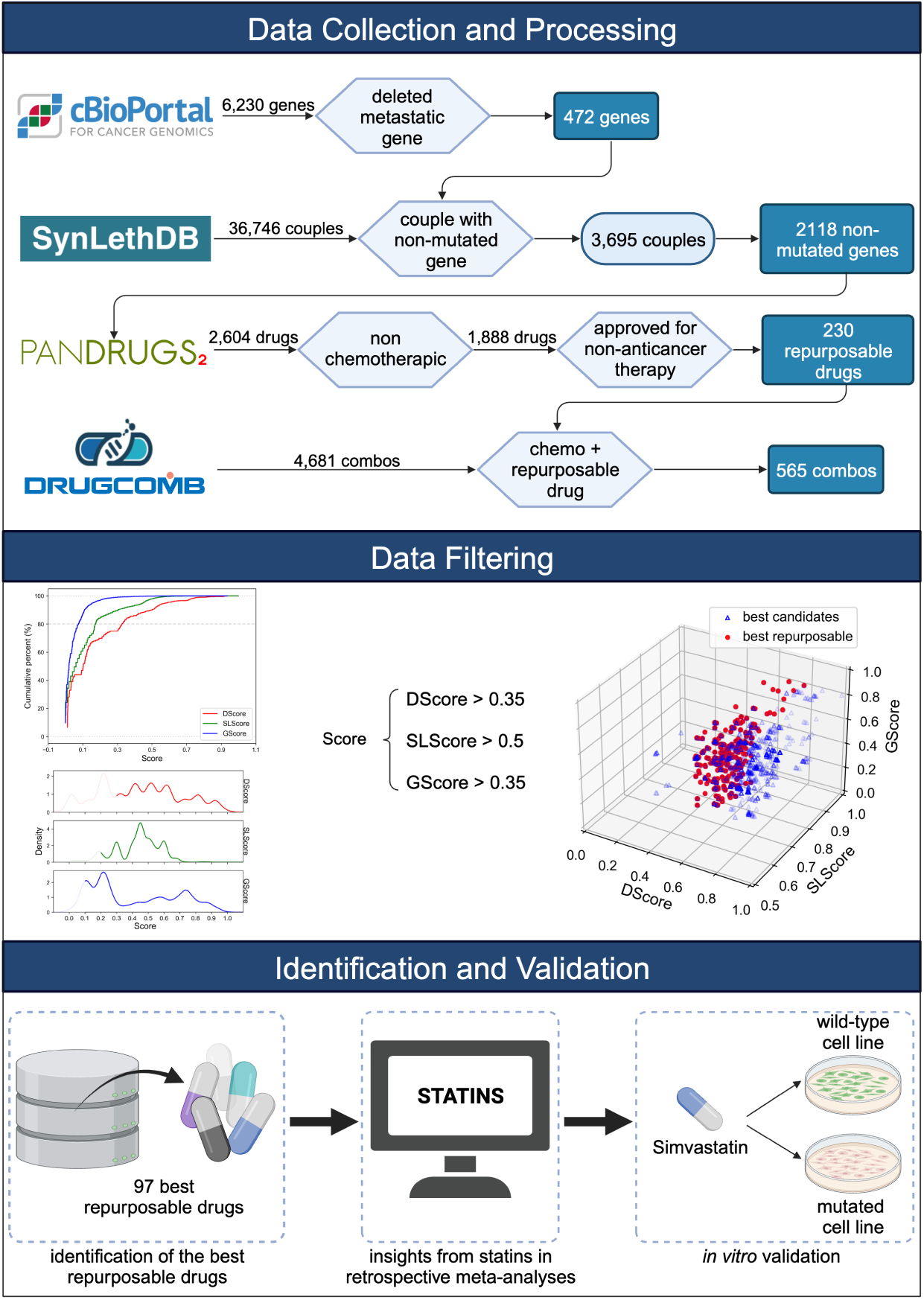
Graphical scheme of the three stages of our research approach: (i) **Data Collection and Processing:** Three types of entities (genes, drugs, and cancer types) are gathered and scrutinized from databases to explore the landscape of migrastatic and anti-metastatic repurposable drugs via SL. The dataset encompasses five types of relationships (metastatic gene, SL gene pair, drug-gene relationship, relationships between drugs and their approved indications, and synergistic drug combination), providing the computational groundwork for drug repositioning. (ii) **Data Filtering**: The dataset is sorted based on GScore, DScore, and SLScore values. The most promising candidates are pinpointed through cumulative percentages and alignment with the density distribution. Optimal scores are identified as GScore *>* 0.35, SLScore *>* 0.5, and DScore *>* 0.35. Within the pool of best candidates, a focused search was conducted to identify those with repurposable potential. (iii) **Identification and Validation**: Among the best repurposable candidates, statins have demonstrated antitumor activity in accordance with retrospective meta-analyses. The *in vitro* experimental validation is carried out with cell lines presenting necessary mutations for susceptibility and wild type cell lines as a negative control.

Synthetic Lethality (SL) pairs with deleted metastatic genes, followed by subsequent *in vitro* validation. Oncogenes associated with metastasis were curated from cBioPortal, resulting in a refined subset of 472 genes. We then sought SL partners of these genes within the SynLethDB database, distilling our initial pool of 36,746 pairs down to a final selection of 3,695 pairs above the fourth quantile, extrapolating 2,118 targetable non-mutated genes. The subsequent phase involved identifying agents that can target these selected non-mutated genes, using associations found within PanDrugs. Our investigation uncovered 2,594 pharmaceutical compounds that target these genes. We focused on non-chemotherapeutic agents, resulting in a selection of 1,888 compounds. Out of these, 230 were repurposable drugs (Fig. 2a). Thus, 18.2%, of the drugs under investigation are currently approved for therapies not related to cancer (Fig. 2b). However, these drugs exhibit potential for repurposing in oncological treatments that leverage the concept of SL.

**Fig. 2:**
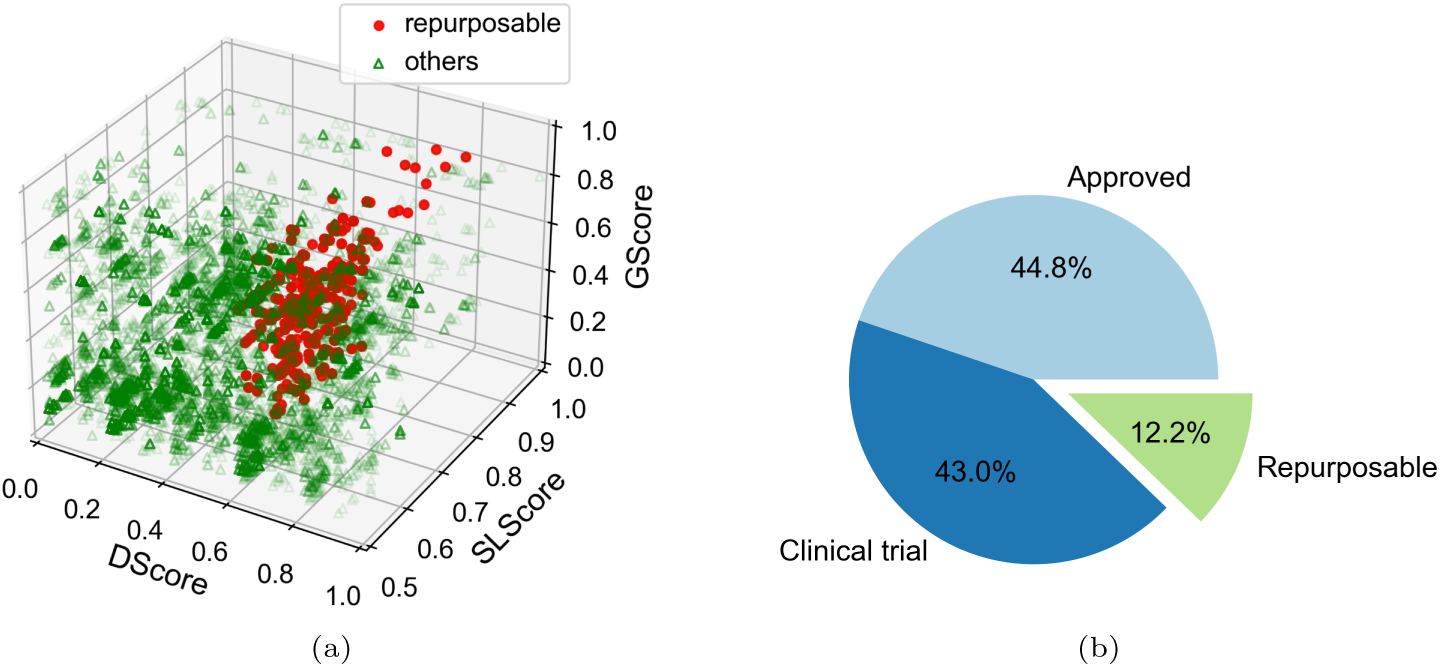
Results of the collection and processing of candidates to identify the repurposable drugs usable for the anticancer Synthetic Lethality approach. **(a)** The three-dimensional representation provides insight into the targeting of non-mutated genes in a Synthetic Lethality association with a deleted metastatic gene. The green triangles signify all the discerned drugs that target non-deleted genes in an SL pair with deleted metastatic genes, and are non-chemotherapeutic, totaling 1,888. Among these, 230 candidates are repurposable (red dot). **(b)** The pie chart provides a visual representation of the distribution of the 1,888 drugs we identified, grouped according to their status. Specifically, 44.8% of these drugs are already approved for cancer treatment, 43.0% are in clinical trial phase, while 12.2% encompass commercially available drugs that are used for non-cancer related treatments but have potential for repurposing in cancer therapy.

We further analysed all potential candidates within our dataset, categorizing them based on their respective GScore, DScore, and SLScore values. Consequently, out of initial candidates, 219 exhibited more compelling characteristics (listed in Supplementary Table 1), and of these, 97 were repurposable. (Fig. 3a, listed in Supplementary Table 2). Notably, our analysis identified approximately 44.3% of the top candidates as potentially repurposable pharmaceuticals (Fig. 3b). This highlights the untapped potential within the existing pharmacopeia and signals a paradigm shift towards repurposing existing compounds for novel therapeutic applications.

**Fig. 3:**
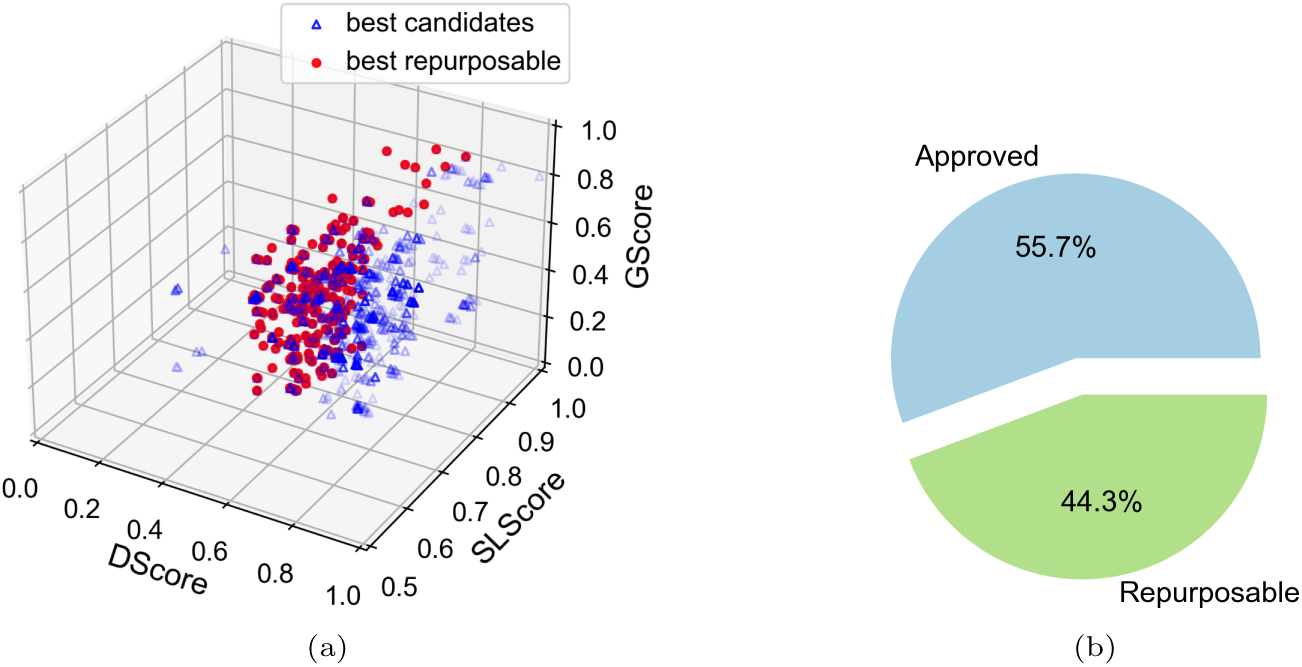
Graphical description of the filtering process applied during the identification of best candidates. **(a)** Through the implementation of statistically determined scoring thresholds (DScore *>* 0.35, SLScore *>* 0.5, GScore *>* 0.35), we have identified a subset of candidates exhibiting superior promise. This has resulted in a pool of 219 best candidates, represented by blue triangles. Within this pool, the number of repurposable drugs is further refined to 97, distinctly visualized as red dots. **(b)** Among the 219 best candidates, 55.7% have already received approval for use in cancer treatment, while 44.3% are commercially available drugs currently used for non-cancer treatments but show potential for repurposing in cancer therapy.

**Table 1:**
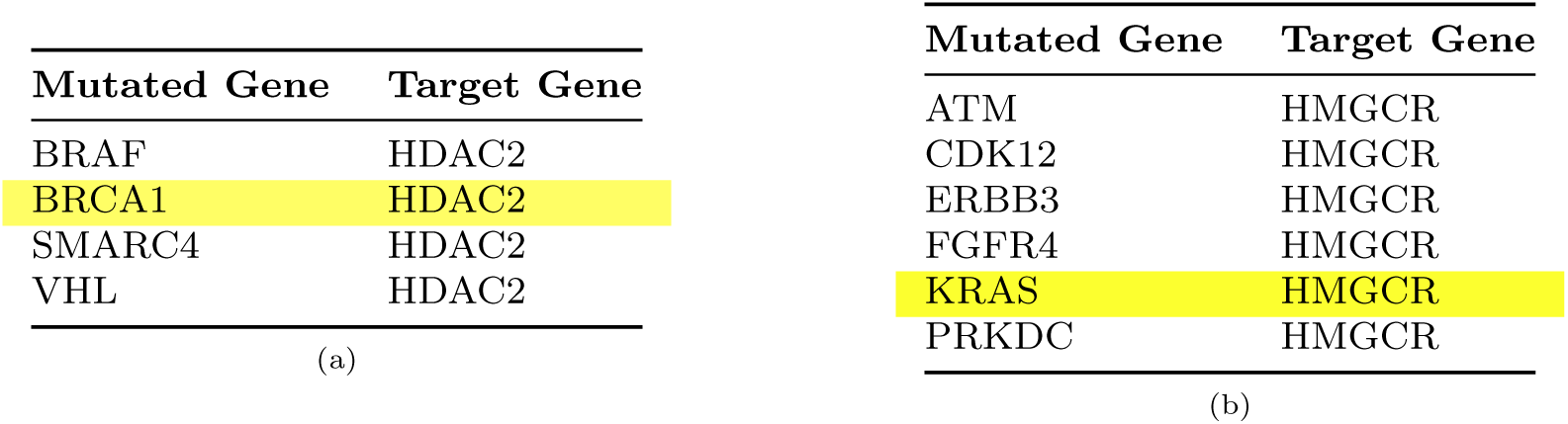
The two tables show the computational findings pertaining to the **(a)** HDAC2 and **(b)** HMGCR genes, and their corresponding Synthetic Lethality pairments with deleted metastatic genes, which are inhibited by drugs from the statin family (Atorvastatin, Lovastatin, Simvastatin, Pravastatin). The SL pairs that we selected for experimental validation are highlighted.

Among the repurpusable pharmaceutical agents that exploit the SL principle, we made thoughtful selections for subsequent experimental validation of data-driven results. Statins emerged as the most compelling options due to the presence in literature of retrospective meta-analyses and experiments that demonstrate their antitumor activity without being able to explain the therapeutic principle beyond this activity [24, 26–29]. Statins target genes HDAC2 and HMGCR which form SL pairs with BRCA1 and KRAS respectively (Tab. 1). Among the statins family identified through computational analysis, we chose Simvastatin as the most clinically relevant candidate for experimental testing.

As an extension to our primary focus, we also conducted an inquiry to identify chemotherapeutic agents that could synergize with Simvastatin. This inquiry is deemed essential, as the integration with a chemotherapeutic agent could potentially facilitate the rapid deployment of the repurposable drug identified here in a clinical setting. From a pool of 4,691 potential synergistic drug pairs, we discovered 565 couples consisting of chemotherapeutic agents paired with repurposed drugs. Notably, Simvastatin is paired with the chemotherapic agent Temozolomide, an oral alkylating agent.

### 2.2 *In vitro* validation of Simvastatin as anticancer agent

Simvastatin, belonging to the statin family widely utilized in the management of cholesterol levels and cardiovascular diseases [30], has emerged as a prime candidate for experimental validation, from our computational analysis. This selection is underscored by retrospective meta-analyses on large patient cohorts that have revealed an unexpected antitumor potential of statins [24, 31]. We speculated that cell lines possessing Synthetic Lethality (SL) pair mutations with genes susceptible to Simvastatin, may possess a heightened sensitivity to this drug (Tab. 2). On the other hand, cell lines devoid of these mutations are anticipated to exhibit resistance to drug treatment. To validate this hypothesis, we performed drug tests to determine the half-maximal inhibitory concentration (IC50) of Simvastatin for several cell lines (MDA-MB-231, HCC1937, OVPA8 and HeLa; refer to Tab. 3). Cell lines harboring mutations susceptible to Simvastatin demonstrated a significant drug response with dosages varying in the range of tens of µM (from 0.01 to 40 µM)(Fig. 4). Specifically, the IC50 was equal to 2.108 *±* 1.11 µM for MDA-MB-231 breast cancer-derived cells, 20.94 *±* 1.07 µM for HCC1937 breast cancer cells, 8 *±* 1.09 µM OVPA8 ovarian cancer cell line. In contrast, coherently with our hypothesis, HeLa cells, which do not carry mutations in the SL genes defined here, did not exhibit any susceptibility, although the drug may have a cytotoxic effect at exposure times longer than 72 hours and/or at higher concentrations. Thus, the experimental data validated our hypothesis, firmly establishing Simvastatin as an antitumor agent, of which the activity is unequivocally due to the phenomenon of SL.

**Fig. 4:**
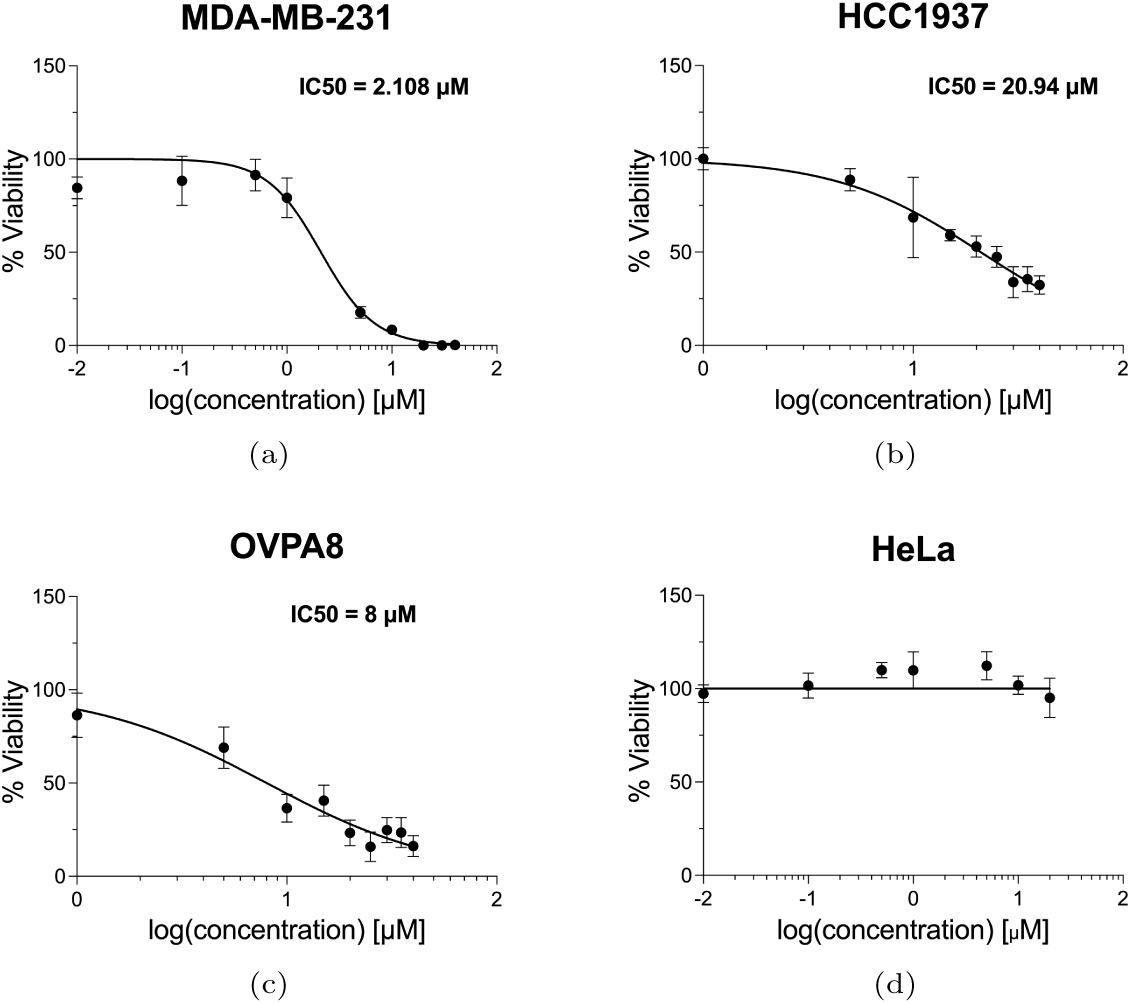
Drug validation of the Synthetic Lethality hypothesis. Half-maximal inhibitory concentration (IC50) is determined for different cancer cell lines, after 72h of treatment with Simvastatin. **(a)** The MDA-MB-231 cell line, harboring a mutated KRAS gene that forms an SL pair with HMGCR, demonstrated an IC50 value of 2.108 µM, R^2^ = 0.9554. **(b)** The HCC1937 cell line, harboring a mutated BRCA1 gene that forms an SL pair with HDAC2, demonstrated an IC50 value of 20.94 µM, R^2^ = 0.8715. **(c)** The OVPA8 cell line, harboring a mutated BRCA1 gene that forms an SL pair with HDAC2, demonstrated an IC50 value of 8 µM, R^2^ = 0.8427. **(d)** The HeLa cell line, devoid of mutations that form SL pairs with the target genes of Simvastatin, remains unaffected across all tested Simvastatin concentrations.

**Table 2:** Cell lines harboring genes that are mutated and in Synthetic Lethality pairs with genes inhibited by Simvastatin, exhibit sensitivity to this drug. In contrast, in the absence of these specific mutations, as in HeLa cells, Simvastatin did not exert any effect. The table reports the specific mutated genes for each cell lines treated with Simvastatin, along with the corresponding IC50 values identified.

| Cancer type | Cell line | Mutated Gene | Target Gene | IC<sub>50</sub> (μM) |
| --- | --- | --- | --- | --- |
| Breast | MDA-MB-231 | KRAS | HMGCR | 2.108 ± 1.11 |
| Breast | HCC1937 | BRCA1 | HDAC2 | 20.94 ± 1.07 |
| Ovarian | OVPA8 | BRCA1 | HDAC2 | 8 ± 1.09 |
| Cervix | HeLa | / | / | no effect |

**Table 3:** Overview of the cell types utilized in the experimental validation and their corresponding gene conditions related to the pharmacological test of Simvastatin (wt: Wild-Type, mut: Mutated).

| MDA-MB-231 | HeLa | HCC1937 | OVPA8 |
| --- | --- | --- | --- |
| BRCA1-wt | BRCA1-wt | BRCA1-mut | BRCA1-mut |
| KRAS-mut | KRAS-wt | KRAS-wt | KRAS-wt |

## 3 Discussion

Computational methodologies are progressively being embraced in the scientific community as a valuable adjunct to traditional experimental procedures, with the objective of expediting discovery processes and unveiling novel targets [32]. Several groups have embarked on computational explorations to pinpoint gene pairs demonstrating Synthetic Lethality (SL) [12, 33], with a subset harnessing the potential of machine learning to uncover new SL couples [34]. Simultaneously, others have delved into databases in pursuit of drugs amenable to repurposing [19]. Yet, no one has proposed a data-driven approach akin to ours, where we have identified repurposable drugs that target genes exhibiting SL when coupled with deleted metastatic genes. This strategy may lead to new antimetastatic and migrastatic therapeutic approach, surpassing the limitations of conventional treatments. Our computational approach, integrating various databases, allowed us to achieve results absent in the literature to date, such as the explanation of the antitumor activity of hydrophobic statins. A review of the existing literature on statin experimentation reveals a multitude of secondary and pleiotropic effects identified by various researchers [27–29]. However, the elucidation of a definitive therapeutic principle within these studies remains elusive. Despite this, retrospective meta-analyses have corroborated their effectiveness in preventing and treating both breast and ovarian metastatic tumors [24, 25, 31, 35, 36]. In this work, we elucidate the anticancer mechanism of action of statins based on Synthetic Lethality analysis. Among the statins identified computationally, Simvastatin was selected for experimental validation. Here, we showed the anticancer mechanism of action of Simvastatin based on Synthetic Lethality (SL) analysis (Fig. 5). One of the genes that we identified is HMGCR, responsible for the inhibition of mevalonate pathway and prenylation [37]. Our SL analysis underscores the pivotal role of Simvastatin in inhibiting the mevalonate pathway, extending beyond its classical function in cholesterol biosynthesis. The suppression of this pathway disrupts the production of key intermediates, such as farnesyl pyrophosphate (FPP) and geranylgeranyl pyrophosphate (GGPP) [38]. This disruption has profound implications for cancer cells that rely heavily on prenylation for the proper functioning of oncogenic proteins [39]. Moreover, our study elucidates how Simvastatin-induced inhibition of protein prenylation acts synergistically with other cancer-related genetic alterations, primarily KRAS, leading to SL. Cancer cells, already harboring mutations that render them susceptible to this disruption, experience a lethal combination of compromised cellular processes, ultimately contributing to their demise. One of the most striking observations in our study is the selective induction of apoptosis in cancer cells treated with Simvastatin. The interconnectedness of disrupted mevalonate pathway intermediates altered protein prenylation, and downstream signaling cascades converges on the activation of apoptotic pathways. The SL induced by Simvastatin represents a finely tuned molecular strategy, ensuring the preferential elimination of cancer cells while sparing normal, healthy cells [12]. Furthermore, the study delves into the specific apoptotic pathways activated by Simvastatin, shedding light on potential targets for future therapeutic interventions. Understanding the intricacies of apoptosis induction in the context of SL provides a roadmap for developing targeted therapies that exploit these vulnerabilities in cancer cells [11]. The SL analysis has unveiled an intriguing interaction between BRCA1 and HDAC2, offering a novel perspective on the potential anticancer mechanism of Simvastatin. BRCA1, a tumor suppressor gene implicated in DNA repair processes [40], is frequently mutated in various cancers, particularly breast and ovarian cancers [41]. Our study indicates that Simvastatin induces SL in cancer cells with BRCA1 mutations when coupled with the inhibition of HDAC2, a histone deacetylase involved in epigenetic regulation [42].

**Fig. 5:**
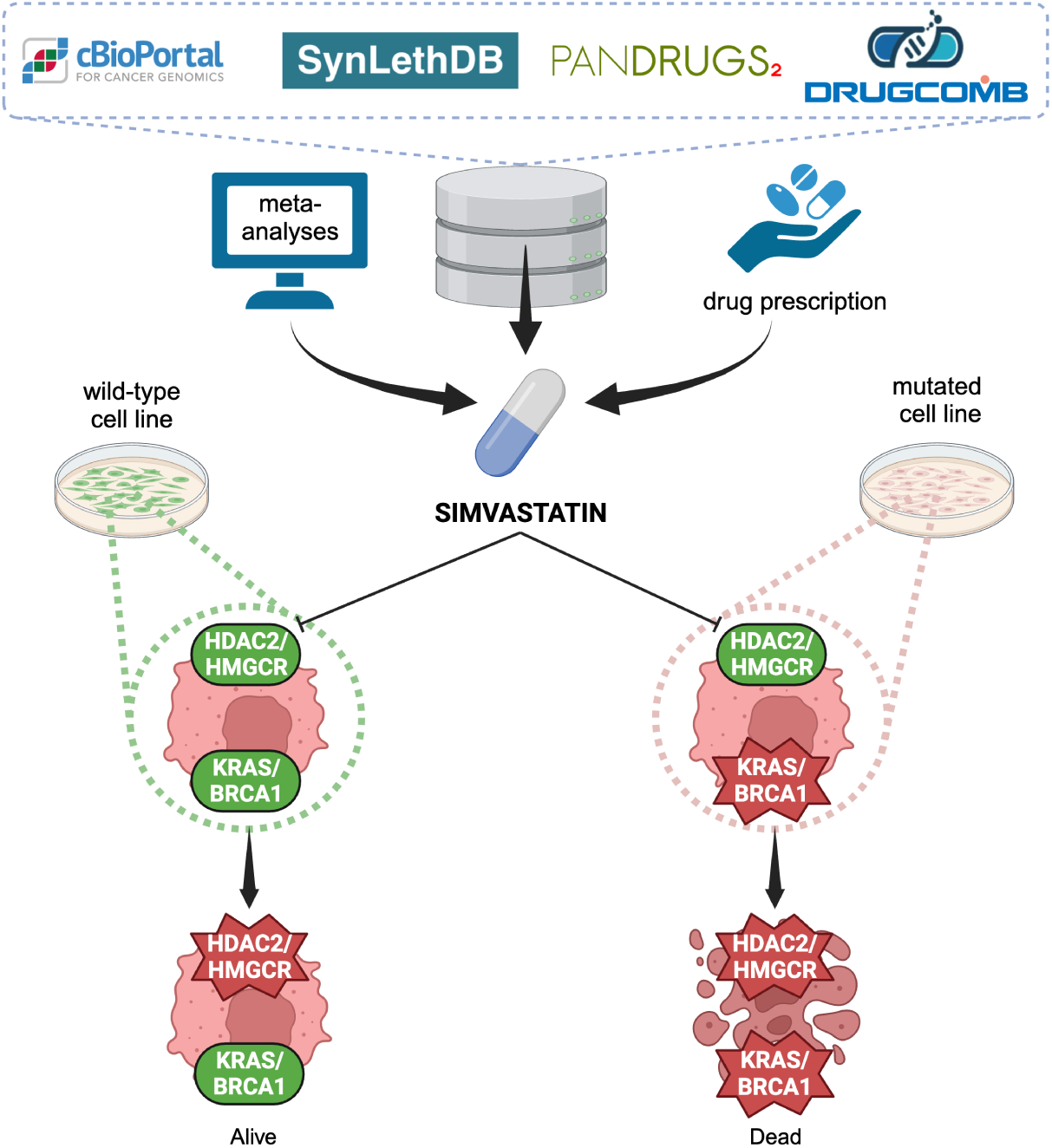
Simvastatin, selected from our data-driven analysis and substantiated by retrospective meta-analyses and extensive current prescriptions data, has demonstrated its antitumor activity rooted in Synthetic Lethality through experimental validation. This compelling evidence irrefutably establishes its therapeutic principle, marking a significant stride in the realm of personalized medicine based on data-driven repurposed drugs.

The connection between BRCA1 and HDAC2 suggests a delicate balance in maintaining genomic integrity. Simvastatin, by targeting HDAC2 in BRCA1-deficient cells, exacerbates the underlying genomic instability, resulting in SL. This interaction is of particular significance in the context of BRCA1-mutated tumors, offering a potential avenue for targeted therapy with Simvastatin. The study’s findings underscore the importance of understanding the intricate molecular relationships within cancer cells, providing a foundation for the development of precision therapies. Simvastatin’s ability to exploit the SL between BRCA1 and HDAC2 may pave the way for innovative treatment strategies for patients with BRCA1-mutated cancers. The SL analysis has identified a compelling interaction between KRAS and HMGCR, unraveling a potential vulnerability that could be leveraged by Simvastatin. KRAS, a proto-oncogene frequently mutated in various cancers, drives uncontrolled cell proliferation and survival [43]. Our study indicates that Simvastatin induces SL when coupled with the inhibition of HMGCR, a key enzyme in the mevalonate pathway. The observed SL suggests that Simvastatin disrupts critical cellular processes in KRAS-mutated cells, further sensitizing them to the inhibition of HMGCR. This interaction holds significance for cancers driven by KRAS mutations, providing a potential avenue for targeted therapy with Simvastatin. The mevalonate pathway, targeted by Simvastatin, plays a crucial role in the survival of KRAS-mutated cells, making this gene couple particularly promising for therapeutic intervention. The study opens up possibilities for the development of tailored treatment strategies for KRAS-mutated cancers. Simvastatin, with its established safety profile, could be repurposed to exploit the SL between KRAS and HMGCR, offering a targeted and potentially more effective approach for patients with limited treatment options.

The identification of SL interactions involving BRCA1-HDAC2 and KRAS-HMGCR couples has substantial clinical implications. Simvastatin, a widely used and well-tolerated medication, may find new therapeutic applications in specific subsets of cancer patients based on their genetic profile. The study emphasizes the importance of patient stratification for optimal treatment outcomes. Clinical trials focusing on these specific gene couples are warranted to validate the safety and efficacy of Simvastatin in the context of SL. Moreover, molecular profiling of tumors to identify patients with BRCA1 mutations or KRAS mutations will be essential for refining treatment strategies and maximizing the potential benefits of Simvastatin-based therapies. Further investigation into the specific molecular mechanisms underlying the SL interactions between BRCA1-HDAC2 and KRAS-HMGCR will deepen our understanding of Simvastatin’s effects on these pathways. Preclinical models and additional genomic analyses can validate and expand upon these findings, providing a robust foundation for clinical translation. Future studies should explore the potential of combination therapies that leverage Simvastatin alongside other targeted agents to enhance the SK observed in specific genetic contexts. This approach may open new possibilities for precision medicine in cancer treatment, addressing the complexity of genetic alterations in a more nuanced manner.

## 4 Conclusion

In conclusion, we integrated extensive data analysis to identify the most promising candidates among drugs that target non-mutated genes involved in Synthetic Lethality (SL) pairs with deleted metastatic genes. The SL analysis sorted out the BRCA1-HDAC2 and KRAS-HMGCR couples, providing valuable insights into potential vulnerabilities in cancer cells that could be targeted by Simvastatin. These findings pave the way for personalized treatment strategies, offering hope for improved outcomes in patients with BRCA1-mutated or KRAS-mutated cancers. Rigorous clinical validation and ongoing research will be pivotal in realizing the full potential of repositioning Simvastatin as a targeted therapy in the evolving landscape of precision oncology. The merits of our framework are highlighted by its potential to expedite drug development by utilizing approved drugs, consequently reducing associated costs and risks.

## 5 Methods

### 5.1 Data Collection and Integration

The computational architecture underpinning our research is predicated on three core entities: genes, drugs, and cancer types. These entities are interconnected via five distinct categories of relationships: associations between cancer metastatic phenotype and their corresponding gene conditions, Synthetic Lethal (SL) gene pairs, connections between drugs and their target genes, relationships between drugs and their approved indications, and the potential for synergy between two drugs. This section delineates the databases, the encompassed information and the filtering path that have been employed in the process of identifying the most suitable candidates.

#### 5.1.1 DataBases

1. cBioPortal [44] has been employed for the identification of gene mutations implicated in metastases. Specifically, it has been done within the context of a pan-cancer metastatic solid tumor study [45]. This comprehensive study encompasses whole-exome and -transcriptome sequencing of 500 adult patients with metastatic solid tumors and primary normal pairs of diverse lineage and biopsy sites. Within this rich dataset, genes are validated as oncogenes through comparison with data from OncoKB [46], helping the study to provide valuable insights into the gene’s expression status in metastatic scenarios.
2. SynLethDB [47] houses all SL pairs discovered to date, collated through various investigative techniques. The database is a rich amalgamation of diverse sources, incorporating experimental data from biochemical assays, literature-derived information, publicly available datasets, computational predictions, and knowledge extracted from textual sources.
3. PanDrugs [48] serves as a robust engine for prioritizing anticancer drug treatments based on individual multi-omics data. It is a culmination of data from an impressive 23 primary sources, resulting in a vast repository of 74,087 drug-target associations involving 4,642 genes and 14,659 unique compounds. It provides comprehensive information on both the drug and the drug’s target gene; it also details the status of the drugs, including those in clinical trial, approved as antitumor, and approved for non-antitumor therapy.
4. DrugComb [49] is a comprehensive drug sensitivity data repository and analysis portal. As an open-access, community-driven data portal, it accumulates, standardizes, and harmonizes the results of drug combination screening studies conducted across a diverse array of cancer cell lines. This database was selected to identify potential synergistic pairs composed of chemotherapeutic and repositioned drug.

#### 5.1.2 Scores

Our selection of best candidates was guided by scores integrated into the aforementioned databases:

1. The Synthetic Lethality Score (SLScore) from SynLethDB, a measure of the reliability of SL based on the strength of evidence. A higher score indicates a more robust consensus derived from multiple types of evidence.
2. The Gene Score (GScore) from PanDrugs, ranging from 0 to 1, evaluates the biological relevance of the gene. It considers factors such as its essentiality, vulnerability, relevance in cancer, biological impact, frequency in genetic variation databases, and clinical implications.
3. The Drug Score (DScore) from PanDrug, spanning from −1 to 1, gauges the suitability of the drug based on factors such as drug-cancer type indications, clinical status, gene-drug relationships, support from curated databases, and collective gene impact.
4. The Zero Interaction Potency (ZIP) synergy score, a key metric in DrugComb, provides a quantification of the degree to which the combined influence of two drugs exceeds the aggregate of their independent effects, under the presumption of non-interaction.

#### 5.1.3 Pipeline

Building on the goal of finding a repurposable therapy that leverages the concept of Synthetic Lethality (SL), the inception of our study involved the identification of oncogenes that were deleted in metastatic conditions, achieved by querying the pancancer metastatic solid tumor study via cBioPortal. The gene selection was predicated on two specific criteria: a deleted expressive state in the metastatic phenotype and an oncogenic classification. The former was inferred from the “Variant Type” information, specifically the “DEL” term, indicative of deleted genes. The latter was facilitated by an integrated dataset with OncoKB in the study; the genes initially identified were cross-referenced with this dataset, selecting those that yielded a positive response to the “Is Cancer Gene (OncoKB)” query.

This curated gene pool of deleted oncogenes in metastases was subsequently cross-referenced with SL pairs within the SynLethDB database. To refine the selection process and mitigate the computational load, we considered only those pairs within the fourth quantile or above, revealing pairs with an SLScore exceeding 0.5. Notably, only one gene from each SL pair was required to be part of the previously curated pool. The list of counterpart genes, corresponding to non-mutated genes, was then sought in PanDrugs to identify potential drug targets. Furthermore, we conducted an exhaustive screening of the database to identify pharmaceutical agents suitable

Among the drugs identified, a further selection was made to identify synergistic combinations with a conventional chemotherapeutic. This research is of considerable importance as the combination with a chemotherapeutic protocol could potentially fast-track its implementation in a clinical setting. This was accomplished via DrugComb, identifying previously discarded chemotherapeutics in combination with repurposable drugs exhibiting a ZIP score exceeding 0.

In our quest to identify the most promising candidates, we employed a tripartite scoring system encompassing drug, synthetic lethality, and gene scores, in according with other example in literature [12]. An initial filtering was conducted using a common principle for identifying the threshold of all scores, namely the 80% of the cumulative percentage, a widely accepted benchmark rooted in the Pareto principle or the 80/20 rule [50]. The calculation of cumulative percentages provides valuable insights into score distribution, trend identification, and impact evaluation, highlighting the relative positioning of specific score points.

The preliminary thresholds for selecting the scores corresponding to cumulative percentages above 80% were identified independently for each curve, specifically at 0.3 for the Drug Score (DScore), 0.2 for Synthetic Lethality pairs (SLScore), and 0.1 for the Gene Score (GScore) (Fig. 6). Through this filtering process, the number of candidates is reduced by 35%. However, given the extensive pool of remaining candidates even after the initial filtering, a secondary, more stringent filtering was deemed necessary. This involved identifying significant thresholds based on each specific score’s probability distribution. The probability distribution of scores is a tool in statistics and data analysis, offering valuable insights into variable understanding, hypothesis testing, and outcome prediction. Taking as a starting point the three scores highlighted by the cumulative percentages filtering, we identified and set the second threshold nearby the first significant peaks after the previous minimum scores, representing the highest likelihood of encountering candidates meeting our filtering criteria. The area under the probability density function curve between two points corresponds to the probability of the variable falling within that score interval. Consequently, the final thresholds were determined to be DScore *>* 0.35, SLScore *>* 0.5, and GScore *>* 0.35, effectively narrowing down the pool of potential candidates (Fig. 7). This rigorous, two-tiered approach ensures a comprehensive and precise identification of the most promising candidates. In the process of calculating the minimum scores, we made a conscious decision to leverage the full potential of the databases, hence, in the case of SynLethDB, we did not restrict it to the fourth quartile. This was primarily because such a limitation would not have facilitated an accurate distribution of probabilities and cumulative percentages. In line with this, the limit determined based on the SLScore was found to be equivalent to the score pinpointed at the fourth quartile.

**Fig. 6:**
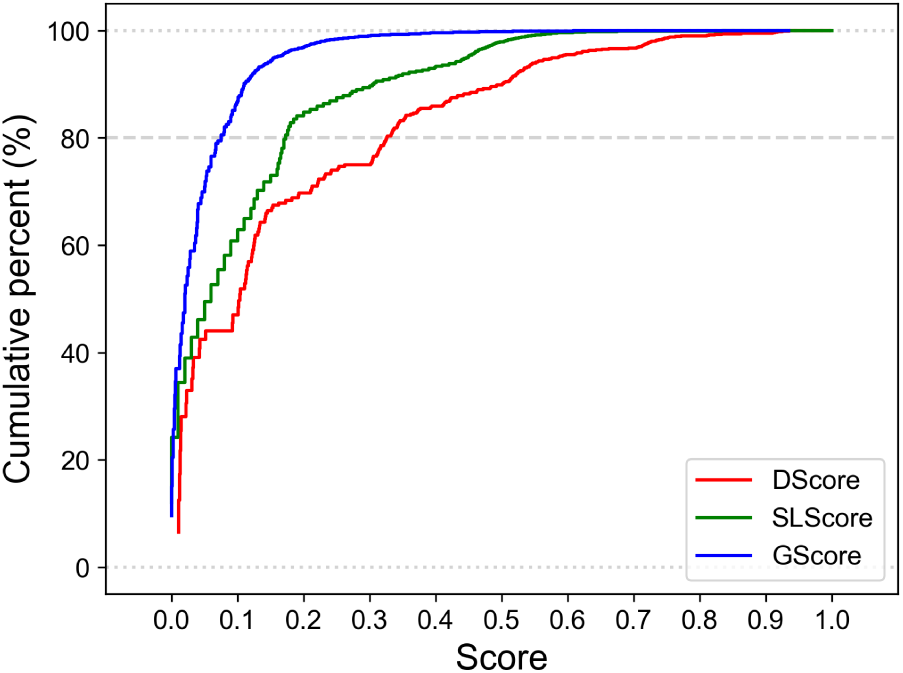
Cumulative distribution curve of the three scores extracted for all possible drugs independently; lines represent the drug score (red), the SL score (green), and the gene score (blue). An 80% threshold has been set to identify the initial minimum significant threshold of scores for candidate filtering, according to Pareto principle. By intersecting the curves and the 80% cutoff, preliminary thresholds were identified at DScore *>* 0.3, SLScore *>* 0.2, GScore *>* 0.1 for repurposing. Since our objective was to develop a comprehensive therapy that combines a repurposed anti-metastatic drug with a conventional cytostatic agent, a first selection was performed by excluding chemotherapeutics as indicated by the “Therapy” entry. Hence, to finally identify repurposable drugs, we utilized the information obtained from PanDrugs in “Status Description”: drugs that presented the term “Cancer Clinical Trials and approved for other pathologies” were considered repurposable.

**Fig. 7:**
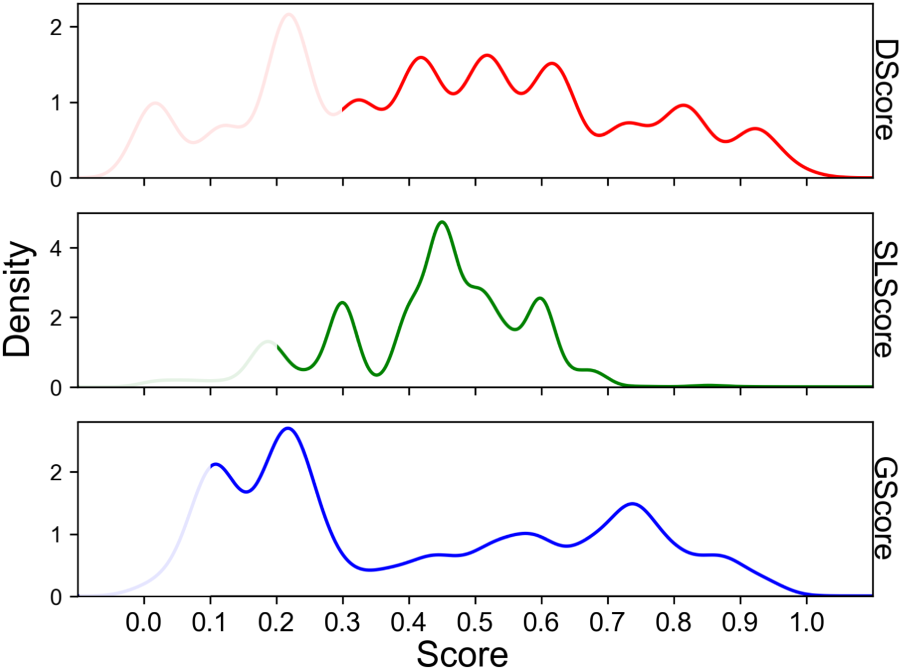
Probability distribution curves for the three scores are delineated; lines signify the drug score (red), the SL score (green), and the gene score (blue). The probability distribution of scores is a tool in statistics and data analysis, offering valuable insights into variable understanding, hypothesis testing, and outcome prediction. The area under the probability density function curve between two points corresponds to the probability that the variable falls within that interval of score. The score corresponding to the first notable peak beyond the minimum threshold, determined by the individual score corresponding to a cumulative percent greater than 80%, is chosen for each score. Consequently, the final thresholds were determined to be DScore *>* 0.35, SLScore *>* 0.5, and GScore *>* 0.35

### 5.2 Cell Cultures & Reagents

#### 5.2.1 Cell Lines, Culture, and Reagents

The MDA-MB-231 and HeLa cell (ATCC, US), a breast adenocarcinoma cell line and a cervical cancer respectively, were maintained in Dulbecco’s Modified Eagle Medium (DMEM, Euroclone, Italy). The HCC1937 (CLS, Germany), a breast ductal carcinoma cell line, and OVPA8 (DSMZ, Germany), a high-grade serous ovarian adenocarcinoma, were cultured in RPMI 1640 medium (Euroclone, Italy). Both media were supplemented with 10% fetal bovine serum (FBS, Euroclone, Italy), 1% l-glutamine (Euroclone, Italy), and 1% penicillin-streptomycin (Euroclone, Italy). All cell lines were incubated at 37°C in a humidified atmosphere containing 5% CO2. Table 3 provides a comprehensive catalogue and elucidation of the cell lines, specifically focusing on their respective states of gene expression of interest. Simvastatin (MK733, MedChemExpress, US), a member of the lipophilic group of statins, was prepared as drug solutions in Dimethyl sulfoxide (DMSO, PanReac AppliChem, Italy) and stored at −80°C.

#### 5.2.2 Cell Viability Assay

To assess the efficacy of Simvastatin on cancer cell lines with various mutations, we designed experiments to draw viability curves and determine the Half-maximal inhibitory concentration (IC50) for each drug on each cell line. The Cell Counting Kit-8 (CCK8, Prodotti Gianni, Italy) assay, a reliable method for gauging cellular metabolic activity as a proxy for cell viability, was employed. Initially, 2,000 cells per well were seeded in 96-well plates and incubated overnight in 100 µL of medium. Subsequently, cells were exposed to a gradient of drug concentrations, ranging from 0.01 to 40 µM, for 72 hours. Notably, the volume of drug diluted in DMSO added to the cell culture was consistently kept below 0.05% in each experiment to minimize its potential influence on the cellular response to the drugs. Post-experiment, the CCK8 was introduced at a 1:10 dilution in culture medium, and incubated for 1-3 hours at 37°C until the solution turned orange. Absorbance at 450 nm was then measured using a Tecan (Infinite M200PRO, Switzerland) microplate reader.

#### 5.2.3 Statistical Analysis

Cell viability data to determine the IC50 were analyzed using GraphPad Prism (Version 9.5.1). The data underwent normalization, with cells treated solely with DMSO established as the benchmark for 100% viability, and wells containing only medium and CCK8 designated as the 0% reference point. The concentrations were converted into logarithmic form to facilitate the construction of the viability curve. This was accomplished by generating a non-linear regression curve that provided an optimal fit with the data, thereby enabling a robust analysis of cell viability across varying drug concentrations. For the appraisal of the congruence between the regression curves and the empirical data, only those instances where the R^2^ values greater than 0.8 were factored into the analysis. Each experimental procedure was executed with a baseline of four replicates. The outcomes are articulated as mean values, augmented by their respective standard deviations.

## Supplementary information

Supplementary Table 1: the table, a result of the integration and synthesis of heterogeneous datasets, is subjected to a filtering procedure based on the corresponding scores. Consequently, it highlights the candidates with the highest potential. Supplementary Table 2: table of the best repurposable candidates.

## Declarations

- Funding: European Research Council (ERC, project BEACONSANDEGG, G.A. 101053122).
- Conflict of interest/Competing interests: : all the authors declare no conflicts of interest/Competing interests.
- Ethics approval: Not applicable.
- Consent to participate: Not applicable.
- Consent for publication: all authors declare consent for publication.
- Availability of data and materials: data are available under reasonable request.
- Code availability: Not applicable.
- Authors’ contributions: GB: computational analysis, experiments, manuscript writing; CT: computational analysis, experiments, manuscript writing; EJ: study design, supervision of experimental analysis; manuscript revision; PP: supervision of computational analysis, manuscript writing; SCarelli: study design, supervision of computational analysis, supervision of experiments, manuscript revision; SCeri: study design, supervision of computational analysis; manuscript revision, fund raising; MTR: study design, supervision of experiments, manuscript revision, fund raising.

## Notes

### Competing Interest Statement

The authors have declared no competing interest.

## References

[1] Siegel, R.L., Miller, K.D., Fuchs, H.E., Jemal, A.: Cancer statistics, 2022. CA: A 17 Cancer Journal for Clinicians 72(1), 7–33 (2022) 10.3322/caac.21708

[2] Horowitz, M., Neeman, E., Sharon, E., Ben-Eliyahu, S.: Exploiting the critical perioperative period to improve long-term cancer outcomes. Nature Reviews Clinical Oncology 12(4), 213–226 (2015) 10.1038/nrclinonc.2014.224

[3] Koval, L., Proshkina, E., Shaposhnikov, M., Moskalev, A.: The role of DNA repair genes in radiation-induced adaptive response in drosophila melanogaster is differential and conditional. Biogerontology 21(1), 45–56 (2019) 10.1007/s10522-019-09842-1

[4] Jabbour, S.K., Kim, S., Haider, S.A., Xu, X., Wu, A., Surakanti, S., Aisner, J., Langenfeld, J., Yue, N.J., Haffty, B.G., Zou, W.: Reduction in tumor volume by cone beam computed tomography predicts overall survival in non-small cell lung cancer treated with chemoradiation therapy. International Journal of Radiation Oncology Biology Physics 92(3), 627–633 (2015) 10.1016/j.ijrobp.2015.02.017

[5] Corrie, P.G.: Cytotoxic chemotherapy: clinical aspects. Medicine 36(1), 24–28 (2008) 10.1016/j.mpmed.2007.10.012

[6] Kilmister, E.J., Koh, S.P., Weth, F.R., Gray, C., Tan, S.T.: Cancer metastasis and treatment resistance: Mechanistic insights and therapeutic targeting of cancer stem cells and the tumor microenvironment. Biomedicines 10(11), 2988 (2022) 10.3390/biomedicines10112988

[7] Olivier, T., Haslam, A., Prasad, V.: Anticancer drugs approved by the us food and drug administration from 2009 to 2020 according to their mechanism of action. JAMA Network Open 4(12), 2138793 (2021) 10.1001/jamanetworkopen.2021.38793

[8] Hadiloo, K., Tahmasebi, S., Esmaeilzadeh, A.: CAR-NKT cell therapy: a new promising paradigm of cancer immunotherapy. Cancer Cell International 23(1) (2023) 10.1186/s12935-023-02923-9

[9] Maalej, K.M., Merhi, M., Inchakalody, V.P., Mestiri, S., Alam, M., Maccalli, C., Cherif, H., Uddin, S., Steinhoff, M., Marincola, F.M., Dermime, S.: CAR-cell therapy in the era of solid tumor treatment: current challenges and emerging therapeutic advances. Molecular Cancer 22(1) (2023) 10.1186/s12943-023-01723-z

[10] Levine, B.L., Pasquini, M.C., Connolly, J.E., Porter, D.L., Gustafson, M.P., Boelens, J.J., Horwitz, E.M., Grupp, S.A., Maus, M.V., Locke, F.L., Ciceri, F., Ruggeri, A., Snowden, J., Heslop, H.E., Mackall, C.L., June, C.H., Sureda, A.M., Perales, M.-A.: Unanswered questions following reports of secondary malignancies after car-t cell therapy. Nature Medicine (2024) 10.1038/s41591-023-02767-w

[11] Topatana, W., Juengpanich, S., Li, S., Cao, J., Hu, J., Lee, J., Suliyanto, K., Ma, D., Zhang, B., Chen, M., Cai, X.: Advances in synthetic lethality for cancer therapy: cellular mechanism and clinical translation. Journal of Hematology & Oncology 13(1) (2020) 10.1186/s13045-020-00956-5

[12] Zhang, B., Tang, C., Yao, Y., Chen, X., Zhou, C., Wei, Z., Xing, F., Chen, L., Cai, X., Zhang, Z., Sun, S., Liu, Q.: The tumor therapy landscape of synthetic lethality. Nature Communications 12(1) (2021) 10.1038/s41467-021-21544-2

[13] Nguyen, B., Fong, C., Luthra, A., Smith, S.A., DiNatale, R.G., Nandakumar, S., Walch, H., Chatila, W.K., Madupuri, R., Kundra, R., et al.: Genomic characterization of metastatic patterns from prospective clinical sequencing of 25,000 patients. Cell 185(3), 563–575 (2022)

[14] Gambardella, V., Tarazona, N., Cejalvo, J.M., Lombardi, P., Huerta, M., Roselló, S., Fleitas, T., Roda, D., Cervantes, A.: Personalized medicine: Recent progress in cancer therapy. Cancers 12(4), 1009 (2020) 10.3390/cancers12041009

[15] O’Neil, N.J., Bailey, M.L., Hieter, P.: Synthetic lethality and cancer. Nature Reviews Genetics 18(10), 613–623 (2017) 10.1038/nrg.2017.47

[16] Kaitin, K.I.: Deconstructing the drug development process: the new face of innovation. Clinical Pharmacology & Therapeutics 87(3), 356–361 (2010) 10.1038/clpt.2009.293

[17] Hua, Y., Dai, X., Xu, Y., Xing, G., Liu, H., Lu, T., Chen, Y., Zhang, Y.: Drug repositioning: Progress and challenges in drug discovery for various diseases. European Journal of Medicinal Chemistry 234, 114239 (2022) 10.1016/j.ejmech.2022.114239

[18] Abdelsayed, M., Kort, E.J., Jovinge, S., Mercola, M.: Repurposing drugs to treat cardiovascular disease in the era of precision medicine. Nature Reviews Cardiology 19(11), 751–764 (2022) 10.1038/s41569-022-00717-6

[19] Santamaría, L.P., Carro, E.U., Uzquiano, M.D., Ruiz, E.M., Gallardo, Y.P., Rodríguez-González, A.: A data-driven methodology towards evaluating the potential of drug repurposing hypotheses. Computational and Structural Biotechnology Journal 19, 4559–4573 (2021)

[20] Flanary, V.L., Fisher, J.L., Wilk, E.J., Howton, T.C., Lasseigne, B.N.: Computational advancements in drug repurposing for cancer combination therapy prediction (2023) 10.20944/preprints202305.1637.v1

[21] US Food and Drug Amministration: Drug Approvals and Databases (2022). https://www.fda.gov/drugs/development-approval-process-drugs/drug-approvals-and-databases

[22] Schein, C.H.: Repurposing approved drugs for cancer therapy. British Medical Bulletin 137(1), 13–27 (2021) 10.1093/bmb/ldaa045

[23] Urquhart, L.: Top drugs and companies by sales in 2018. Nature Reviews Drug Discovery (2019) 10.1038/d41573-019-00049-0

[24] Borgquist, S., Broberg, P., Tojjar, J., Olsson, H.: Statin use and breast cancer survival – a swedish nationwide study. BMC Cancer 19(1) (2019) 10.1186/s12885-018-5263-z

[25] Wang, Q., Zhi, Z., Han, H., Zhao, Q., Wang, X., Cao, S., Zhao, J.: Statin use improves the prognosis of ovarian cancer: An updated and comprehensive meta-analysis. Oncology Letters 25(2) (2022) 10.3892/ol.2022.13648

[26] Cafforio, P., Dammacco, F., Gernone, A., Silvestris, F.: Statins activate the mitochondrial pathway of apoptosis in human lymphoblasts and myeloma cells. Carcinogenesis 26(5), 883–891 (2005) 10.1093/carcin/bgi036

[27] Zahedipour, F., Butler, A.E., Rizzo, M., Sahebkar, A.: Statins and angiogenesis in non-cardiovascular diseases. Drug Discovery Today 27(10), 103320 (2022) 10.1016/j.drudis.2022.07.005

[28] Jin, H., He, Y., Zhao, P., Hu, Y., Tao, J., Chen, J., Huang, Y.: Targeting lipid metabolism to overcome emt-associated drug resistance via integrin *β*3/fak pathway and tumor-associated macrophage repolarization using legumain-activatable delivery. Theranostics 9(1), 265–278 (2019) 10.7150/thno.27246

[29] Guerrab, A.E., Bamdad, M., Kwiatkowski, F., Bignon, Y.-J., Penault-Llorca, F., Aubel, C.: Anti-egfr monoclonal antibodies and egfr tyrosine kinase inhibitors as combination therapy for triple-negative breast cancer. Oncotarget 7(45), 73618– 73637 (2016) 10.18632/oncotarget.12037

[30] Pedersen, T.R., Tobert, J.A.: Simvastatin: a review. Expert Opinion on Pharmacotherapy 5(12), 2583–2596 (2004) 10.1517/14656566.5.12.2583

[31] Inasu, M., Feldt, M., Jernström, H., Borgquist, S., Harborg, S.: Statin use and patterns of breast cancer recurrence in the malmö diet and cancer study. The Breast 61, 123–128 (2022) 10.1016/j.breast.2022.01.003

[32] Nature Methods 18(7), 695–695 (2021) 10.1038/s41592-021-01215-2

[33] Schäffer, A.A., Chung, Y., Kammula, A.V., Ruppin, E., Lee, J.S.: A systematic analysis of the landscape of synthetic lethality-driven precision oncology. Med 5(1), 73–89 (2024)

[34] Testa, C., Pidò, S., Jacchetti, E., Raimondi, M.T., Ceri, S., Pinoli, P.: Inference of synthetically lethal pairs of genes involved in metastatic processes via non-negative matrix tri-factorization. In: Proceedings of the 2023 15th International Conference on Bioinformatics and Biomedical Technology, pp. 47–53 (2023)

[35] Majidi, A., Na, R., Jordan, S.J., De Fazio, A., Webb, P.M.: Statin use and survival following a diagnosis of ovarian cancer: A prospective observational study. International Journal of Cancer 148(7), 1608–1615 (2020) 10.1002/ijc.33333

[36] Couttenier, A., Lacroix, O., Vaes, E., Cardwell, C.R., De Schutter, H., Robert, A.: Statin use is associated with improved survival in ovarian cancer: A retrospective population-based study. PLOS ONE 12(12), 0189233 (2017) 10.1371/journal.pone.0189233

[37] Goldstein, J.L., Brown, M.S.: Regulation of the mevalonate pathway. Nature 343(6257), 425–430 (1990) 10.1038/343425a0

[38] Manaswiyoungkul, P., Araujo, E.D., Gunning, P.T.: Targeting prenylation inhibition through the mevalonate pathway. RSC Medicinal Chemistry 11(1), 51–71 (2020) 10.1039/c9md00442d

[39] Alannan, M., Trézéguet, V., Amoêdo, N.D., Rossignol, R., Mahfouf, W., Rezvani, H.R., Dittrich-Domergue, F., Moreau, P., Lacomme, S., Gontier, E., Grosset, C.F., Badran, B., Fayyad-Kazan, H., Merched, A.J.: Rewiring lipid metabolism by targeting pcsk9 and hmgcr to treat liver cancer. Cancers 15(1), 3 (2022) 10.3390/cancers15010003

[40] Deng, C.-X.: Roles of brca1 in dna damage repair: a link between development and cancer. Human Molecular Genetics 12(90001), 113–123 (2003) 10.1093/hmg/ddg082

[41] Fu, X., Tan, W., Song, Q., Pei, H., Li, J.: Brca1 and breast cancer: Molecular mechanisms and therapeutic strategies. Frontiers in Cell and Developmental Biology 10 (2022) 10.3389/fcell.2022.813457

[42] Park, S.-Y., Kim, J.-S.: A short guide to histone deacetylases including recent progress on class ii enzymes. Experimental & molecular medicine 52(2), 204–212 (2020)

[43] Zhu, G., Pei, L., Xia, H., Tang, Q., Bi, F.: Role of oncogenic kras in the prognosis, diagnosis and treatment of colorectal cancer. Molecular Cancer 20(1) (2021) 10.1186/s12943-021-01441-4

[44] Cerami, E., Gao, J., Dogrusoz, U., Gross, B.E., Sumer, S.O., Aksoy, B.A., Jacobsen, A., Byrne, C.J., Heuer, M.L., Larsson, E., et al.: The cbio cancer genomics portal: an open platform for exploring multidimensional cancer genomics data. Cancer discovery 2(5), 401–404 (2012)

[45] Robinson, D.R., Wu, Y.-M., Lonigro, R.J., Vats, P., Cobain, E., Everett, J., Cao, X., Rabban, E., Kumar-Sinha, C., Raymond, V., Schuetze, S., Alva, A., Siddiqui, J., Chugh, R., Worden, F., Zalupski, M.M., Innis, J., Mody, R.J., Tomlins, S.A., Lucas, D., Baker, L.H., Ramnath, N., Schott, A.F., Hayes, D.F., Vijai, J., Offit, K., Stoffel, E.M., Roberts, J.S., Smith, D.C., Kunju, L.P., Talpaz, M., Cieślik, M., Chinnaiyan, A.M.: Integrative clinical genomics of metastatic cancer. Nature 548(7667), 297–303 (2017) 10.1038/nature23306

[46] Chakravarty, D., Gao, J., Phillips, S., Kundra, R., Zhang, H., Wang, J., Rudolph, J.E., Yaeger, R., Soumerai, T., Nissan, M.H., et al.: Oncokb: a precision oncology knowledge base. JCO precision oncology 1, 1–16 (2017)

[47] Wang, J., Wu, M., Huang, X., Wang, L., Zhang, S., Liu, H., Zheng, J.: SynLethDB 2.0: a web-based knowledge graph database on synthetic lethality for novel anti-cancer drug discovery. Database 2022 (2022) 10.1093/database/baac030

[48] Piñeiro-Yáñez, E., Reboiro-Jato, M., Gómez-López, G., Perales-Patón, J., Troulé, K., Rodríguez, J.M., Tejero, H., Shimamura, T., López-Casas, P.P., Carretero, J., Valencia, A., Hidalgo, M., Glez-Peña, D., Al-Shahrour, F.: PanDrugs: a novel method to prioritize anticancer drug treatments according to individual genomic data. Genome Medicine 10(1) (2018) 10.1186/s13073-018-0546-1

[49] Zagidullin, B., Aldahdooh, J., Zheng, S., Wang, W., Wang, Y., Saad, J., Malyutina, A., Jafari, M., Tanoli, Z., Pessia, A., Tang, J.: DrugComb: an integrative cancer drug combination data portal. Nucleic Acids Research 47(W1), 43–51 (2019) 10.1093/nar/gkz337

[50] Erridge, P.: The pareto principle. British Dental Journal 201(7), 419–419 (2006) 10.1038/sj.bdj.4814131

